# Genetic drift acts strongly on within-host influenza virus populations during acute infection but does not act alone

**DOI:** 10.1101/2025.08.27.672713

**Authors:** Yike Teresa Shi, Michael A. Martin, Daniel B. Weissman, Katia Koelle

**Affiliations:** Department of Biology, Emory University, Atlanta, GA, USA; Department of Pathology, Johns Hopkins School of Medicine, Baltimore, MD, USA; Department of Epidemiology, Johns Hopkins Bloomberg School of Public Health, Baltimore, MD, USA; Department of Physics, Emory University, Atlanta, GA, USA; Emory Center of Excellence for Influenza Research and Response (CEIRR), Atlanta GA, USA

**Keywords:** intrahost viral evolution, influenza A virus evolution, genetic drift

## Abstract

The evolutionary dynamics of seasonal influenza A viruses (IAVs) have been well characterized at the population level, with antigenic drift known to be a major force in driving strain turnover. The evolution of IAV populations at the within-host level, however, is still less well characterized. Improving our understanding of within-host IAV evolution has the potential to shed light on the source of new strains, including new antigenic variants, at the population level. Existing studies have pointed towards the role that stochastic processes play in shaping within-host viral evolution in acute infections of both humans and pigs. Here, we apply a population genetic model called the ‘Beta-with-Spikes’ approximation to longitudinal intrahost Single Nucleotide Variant (iSNV) frequency data to quantify the extent of genetic drift acting on IAV populations at the within-host scale. We estimate small effective population sizes in both human IAV infections (*N*_E_ = 41, 95% confidence interval: [22-72]) and swine IAV infections (*N*_E_ = 10, 95% confidence interval: [8-14]). Moreover, we evaluate the consistency of the observed iSNV dynamics with Wright-Fisher model simulations. For the human IAV dataset that we analyze, we find that observed within-host IAV evolutionary dynamics are consistent with this classic model at the estimated low effective population size. However, for the swine IAV dataset, we find statistical evidence for rejecting the classic Wright-Fisher model as the only process governing within-host iSNV frequency dynamics. Our results contribute to the growing number of studies that point towards the important role of genetic drift in shaping patterns of genetic diversity in IAV populations within acutely infected hosts. It further raises questions about whether and what other processes, such as spatial compartmentalization, viral progeny production dynamics with strong skew, or selection, may be needed to explain patterns of within-host IAV evolution.

## Introduction

Viral adaptation at the population level ultimately depends on genetic variation that is generated during viral replication at the within-host scale. Several different mechanisms, however, can underlie the path from mutation generation to spread at the population level. One possibility is that advantageous mutations that are generated during replication (e.g., those altering the antigenicity of a virus) are efficiently selected at the within-host scale, and then subsequently spread at the level of the population. Alternatively, selection may be inefficient at the within-host scale, with population-level spread of advantageous mutations occurring largely due to selection at this higher organizational scale. Previous analyses of synonymous and nonsynonymous viral genetic variation within hosts and at the population level have found strong support for the latter mechanism in human influenza viruses, with both purifying selection and positive selection (at antigenic sites) acting more strongly at the population level than at the within-host level (Xue and Bloom, 2020). Consistent with this finding, three earlier within-host studies indicated that IAV diversity was limited in acutely infected individuals, and that the diversity that was observed was largely shaped by genetic drift and purifying selection (Dinis et al., 2016; Debbink et al., 2017; McCrone et al., 2018). Positive selection on within-host IAV populations was not detected in any of these studies. Similar results to these were found for within-host IAV populations in swine hosts experiencing acute infection (VanInsberghe et al., 2024).

Building on the McCrone et al. (2018) study, McCrone et al. (2020) applied a population genetic model to their intrahost Single Nucleotide Variant (iSNV) data to quantify the strength of genetic drift in these within-host IAV populations. Relying on a diffusion approximation, this analysis estimated the effective population size (*N*_E_) of within-host IAV populations in acute humans infections to be very small, on the order of 32-72 virions. A more recent analysis of IAV evolution based on serial samples from 143 acutely infected individuals estimated within-host *N*_E_ values of 176-284 using approximate Bayesian computation (ABC) with Wright-Fisher simulations and further found evidence for positive selection occurring at 9-11% of the variable sites (Bendall et al., 2024). As such, findings from this recent analysis differ to some extent from the previous ones, finding lower levels of genetic drift and more evidence of positive selection. Finally, a study of within-host IAV evolution in young children found evidence for low viral diversity and purifying selection in seasonal H3N2 infections early on in the course of infection and accumulation of nonsynonymous variants starting around 3–4 days post-symptom onset (Han et al., 2021).

Here, similarly to some of the above studies, we aim to quantify the effective population size *N*_E_ of within-host IAV populations, and moreover, ask whether the evolutionary dynamics observed in within-host IAV populations are consistent with the classic Wright-Fisher model of evolution. We first quantify *N*_E_ using McCrone and coauthors’ previously published within-host IAV data from humans (McCrone et al., 2018) as well as previously published within-host IAV data from swine (VanInsberghe et al., 2024). We quantify the extent of genetic drift in these within-host IAV populations, however, using a different population genetic model, namely the ‘Beta-with-Spikes’ model (Tataru et al., 2015). We use this model due to its demonstrated ability to capture the distribution of allele frequencies over time under a Wright-Fisher model with both large and small population sizes. In contrast, the diffusion approximation is only considered a good approximation when effective population sizes are large. Our analyses support very small within-host viral effective population sizes in both acutely infected humans and swine. Furthermore, for the human IAV dataset we analyze, we find that observed within-host IAV evolutionary dynamics are consistent with dynamics arising from the classic Wright-Fisher model. However, for the swine IAV dataset, we instead find that the classic Wright-Fisher model cannot faithfully reproduce observed patterns of iSNV frequency changes. Alternative models, including ones that describe offspring distributions with strong skew (Okada and Hallatschek, 2021) or ones that include spatial compartmentalization (Gallagher et al., 2018), may thus need to be considered to explain observed changes in iSNV frequencies that are observed in these within-host swine IAV populations.

## Methods

### Overview of the Beta-with-Spikes model

We estimate the effective population size *N*_E_ of within-host IAV populations using a population genetic model called the Beta-with-Spikes model (Tataru et al., 2015). This model approximates the distribution of allele frequencies (DAF) that would result from a Wright-Fisher model over discrete generations. As its name implies, the model uses an adjusted form of the beta distribution. The adjusted form of this distribution includes two spikes at frequencies of 0.0 and 1.0 that account for the probabilities of loss and fixation of alleles, respectively. The distribution of allele frequencies under the Beta-with-Spikes model in generation *t* is given by equation (8) in Tataru et al. (2015), reproduced here:

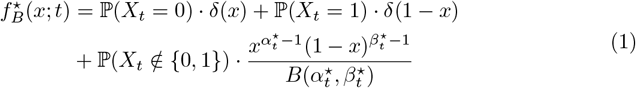

where *δ*(*x*) is the Dirac delta function. The three terms on the left-hand-side of this equation correspond to the probability mass of allele loss, the probability mass of allele fixation, and the probability densities of allele frequencies between (but excluding) 0 and 1, respectively. The calculation for the beta distribution’s shape parameters 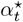 and 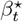 as well as the probabilities for loss and fixation for each generation *t* can be found in Tataru et al. (2015).

To familiarize the reader with the evolutionary dynamics of the Beta-with-Spikes model, we show in Figure 1 simulated DAFs under this model for three different effective population sizes, each starting with the same DAF in generation 0. Figure 1A shows how the DAF changes when the effective population size *N*_E_ is very small (*N*_E_ = 20): the distribution rapidly spreads out from its initial distribution within a small number of generations. By generation 6, allele loss (*X*_*t*_ = 0) already accounts for upwards of 12% of the probability mass. At larger values of *N*_E_ (Figures 1B, 1C), the DAF spreads out from its initial distribution more slowly, as anticipated. For example, when *N*_E_ = 500, allele loss and fixation accounts for less than 1% of the probability mass by generation 6 (Figures 1C).

**Figure 1.**
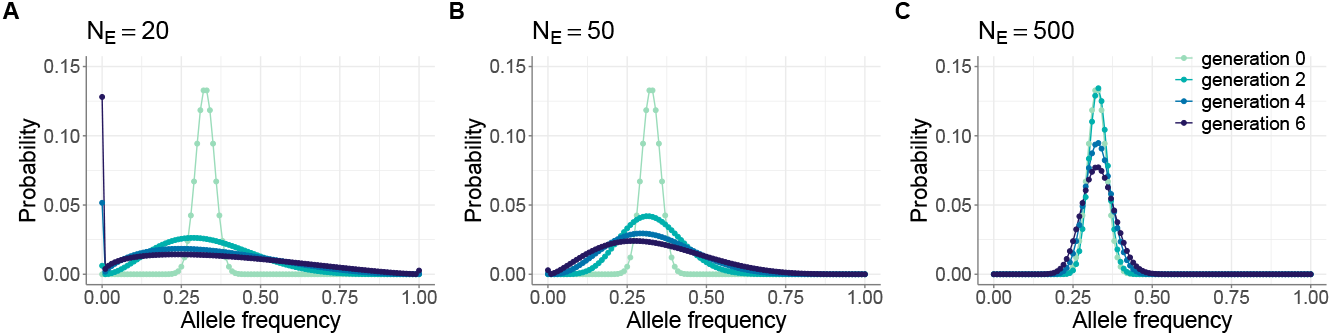
Simulated distribution of allele frequencies (DAFs) under the Beta-with-Spikes approximation for the Wright-Fisher model. Panels (A)-(C) provide simulations under different effective population sizes: (A) *N*_E_ = 20, (B) *N*_E_ = 50, (C) *N*_E_ = 500. In all panels, the initial iSNV frequency distribution in generation 0 was parameterized with a mean value of *p*_0_ = 0.325 and variance given by *v*_0_ = *mp*_0_(1 − *p*_0_), with *m* = 0.004. DAFs from generations 2, 4, and 6 are plotted in each panel to show changes in DAFs across generations. The DAFs in panels (A)-(C) are shown by plotting probability masses in 0.01 frequency intervals.

### Within-host human influenza A virus data

We first analyzed a previously-published human influenza A virus dataset (McCrone et al., 2018), sourced from a community-based cohort study. This dataset includes deep-sequencing data from 43 longitudinally-sampled individuals. Each of these individuals was sampled exactly twice between −2 and 6 days post symptom onset. Variants were called at a minor allele frequency threshold of 2% from sequencing reads accessed through the NCBI Sequencing Read Archive (NCBI SRA BioProject PRJNA412631). For each of these 43 individuals, Table S1 lists the two sample collection dates and the identities and frequencies of all of the identified iSNVs.

We perform two separate analyses, on different subsets of these data. We first estimate *N*_E_ using the subset of iSNVs that were detected above the variant-calling threshold at the first of the two sample collection time points. This includes iSNVs that were still detected above the variant-calling threshold at the second time point as well as iSNVs that were no longer detected or fell below the variant-calling threshold at the second time point. To avoid bias that could result from genetic linkage, we downsample this set of iSNVs to one iSNV per individual by selecting the iSNV that had a frequency closest to 50% at the first time point (and is therefore the most informative of *N*_E_). The downsampled subset of iSNVs comprises 30 paired-time observations. We refer to this dataset as human IAV data subset 1. Table S1 indicates the iSNVs that are included in data subset 1 and Figure 2A plots the frequency dynamics of these data subset 1 iSNVs. Our second analysis estimates *N*_E_ using the subset of iSNVs that were detected above the variant-calling threshold at the second time point but fell below the variant-calling threshold (including those that were undetected) at the first of the two time points. To again avoid bias that could result from genetic linkage, we downsample this set of iSNVs to one per individual, this time selecting the iSNV we keep in our data set at random. The downsampled subset of iSNVs comprises 38 paired-time observations. We refer to this dataset as human IAV data subset 2. Table S1 indicates the iSNVs that are included in data subset 2 and Figure 3A plots the frequency dynamics of these data subset 2 iSNVs.

**Figure 2.**
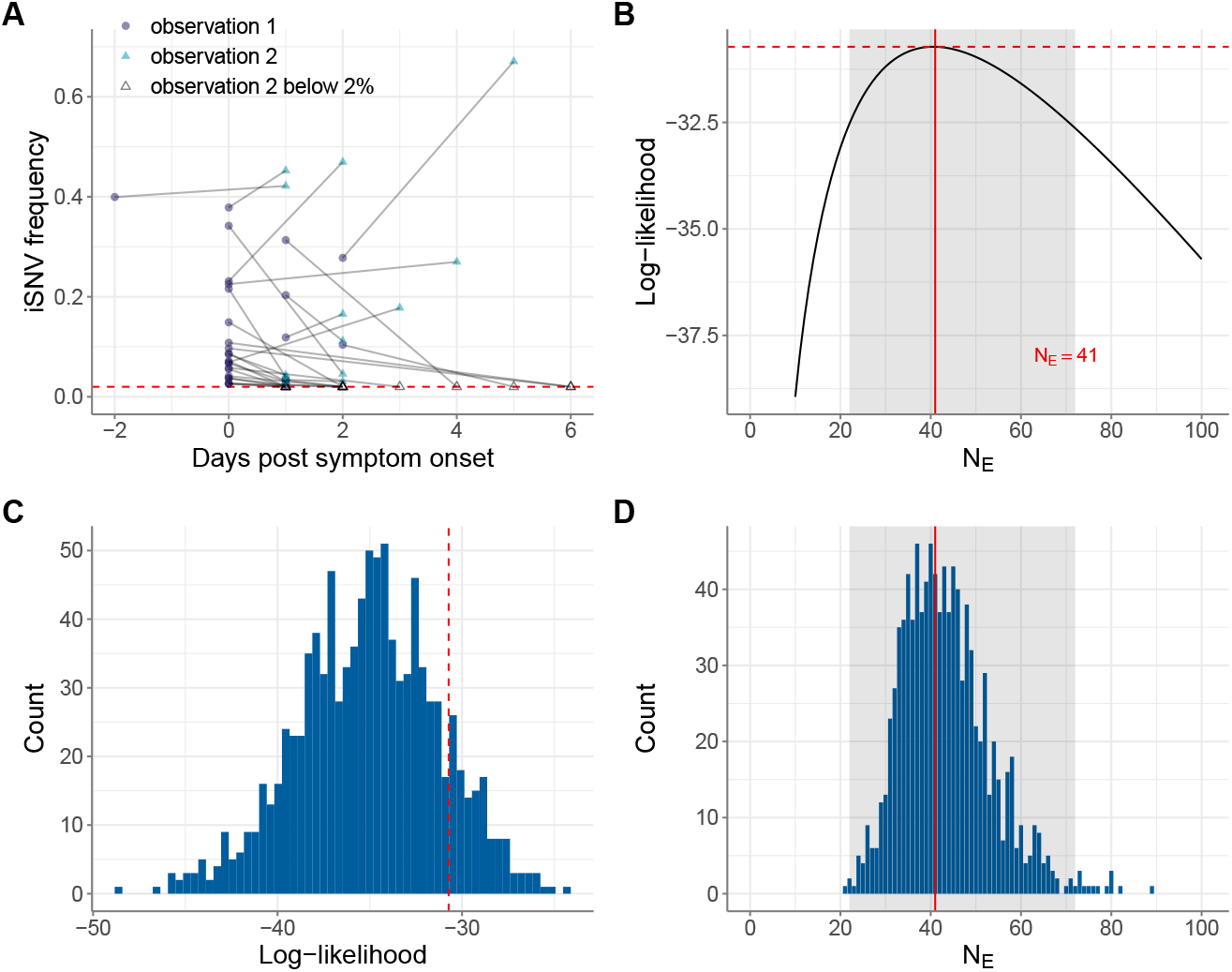
Estimates of within-host IAV effective population size from iSNVs present in the first sampled timepoint (human data subset 1). (A) Allele frequency changes between the first observation time point and the second observation time point. Allele frequencies are plotted by day of symptom onset of the infected individual. The red dashed line shows the variant-calling threshold of 2%. Allele frequencies that fall under this threshold are shown at the threshold. (B) Calculated log-likelihood values across a range of effective population sizes. Solid red line shows the maximum likelihood estimate (MLE) of *N*_E_. Dashed red line shows the log-likelihood value for the MLE of *N*_E_. The shaded region shows the 95% confidence interval around the MLE of *N*_E_. (C) Calculated log-likelihood values for 1000 mock iSNV datasets that were generated by forward simulation of the Beta-with-Spikes model with *N*_E_ = 41 (blue histogram). The log-likelihood value at *N*_E_ = 41 calculated from human data subset 1 is shown with a dashed red line. (D) Maximum likelihood estimates of *N*_E_ from the 1000 simulated datasets, obtained using the Beta-with-Spikes model (blue histogram). The MLE of *N*_E_ = 41 from human data subset 1 is shown with a solid red line. The shaded region shows the 95% confidence interval shown in panel B.

**Figure 3.**
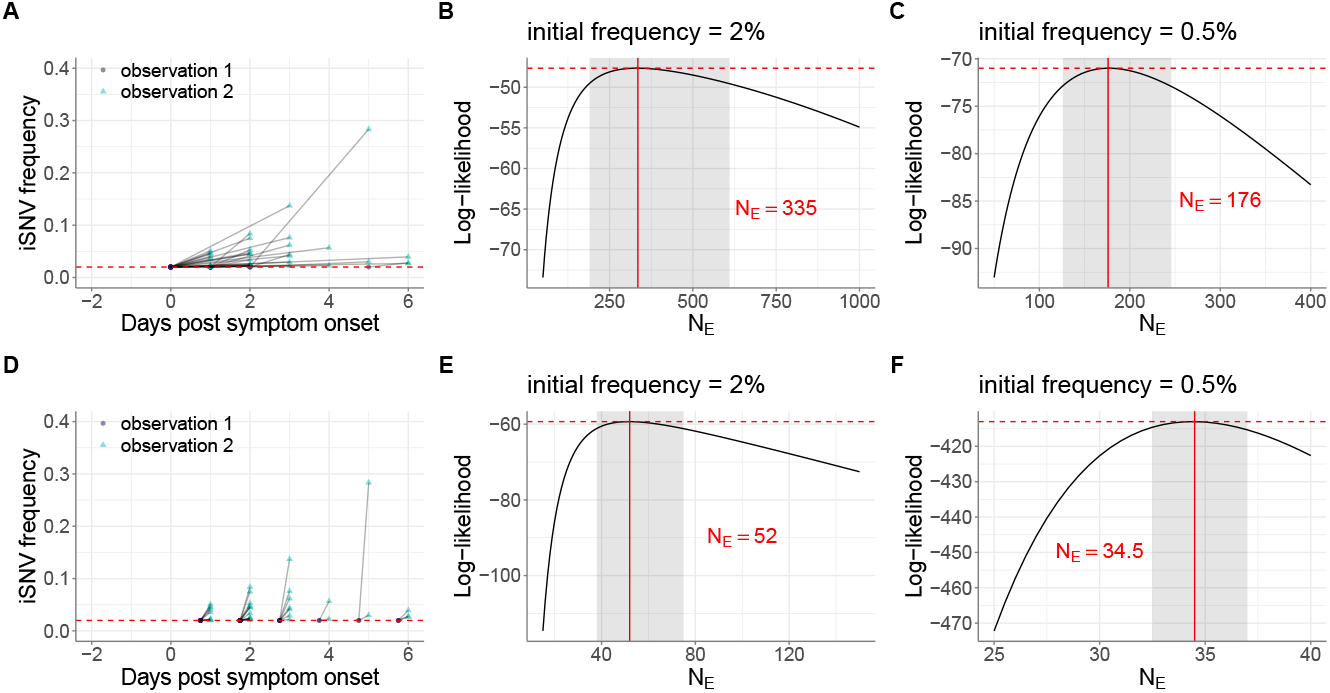
Estimates of within-host IAV effective population sizes from iSNVs present only in the second sampled timepoint (human data subset 2). (A) Allele frequencies at the first and second observation time points in human data subset 2. As in Figure 2A, iSNV frequencies are plotted according to the day of symptom onset and the dashed red line shows the variant-calling threshold of 2%. Below-the-variant-calling-threshold iSNV frequencies at the first time point are shown at 2%. (B) Calculated log-likelihood values across a range of effective population sizes when iSNVs at the first observation time point are assumed to be present at a frequency of 2%. (C) Calculated log-likelihood values across a range of effective population sizes when iSNVs at the first observation time point are assumed to be present at a frequency of 0.5%. (D) Allele frequencies from the paired-time observations in human data subset 2, assuming that the iSNV frequencies were present at the variant-calling threshold a single viral generation prior to the second observation time point. As in Figure 3A, iSNV frequencies are plotted according to the day of symptom onset and the dashed red line shows the variant-calling threshold of 2%. (E) Calculated log-likelihood values across a range of effective population sizes for the modified human data subset 2 shown in panel (D). (F) As in panel (E), calculated log-likelihood values across a range of effective population sizes when iSNVs at the first time point are instead assumed to be present at a frequency of 0.5%. In panels (B), (C), (E), and (F), the solid red lines show the MLE of *N*_E_ and dashed red lines show the log-likelihood value for the MLE of *N*_E_. The shaded regions show the 95% confidence interval around the MLE of *N*_E_.

### Within-host swine influenza A virus data

We further analyzed a previously-published swine influenza A virus dataset (VanInsberghe et al., 2024), sourced from a week-long county fair. This dataset includes deep-sequencing data from 82 longitudinally-sampled pigs with single-subtype infections. Approximately 6% of the sampled pigs have more than 2 longitudinal samples (range: 3-5 samples). Variants were called from sequencing reads accessed through NCBI SRA (BioProject PRJNA1051292) with a minor allele frequency threshold of 2%, as for the human influenza A virus infections.

We generated analogous datasets to human data subset 1 and human data subset 2, using similar approaches as those described above. We refer to these swine datasets as swine data subset 1 and swine data subset 2, respectively. For swine data subset 1, we again downsampled from the set of data subset 1-eligible iSNVs to 1 iSNV per pair of adjacent time points by selecting the iSNV with frequency closest to 50% at the first time point of the pair. For swine data subset 2, we again downsampled the set of iSNVs analyzed to 1 iSNV per pair of adjacent time points by selecting an iSNV from the set of data subset 2-eligible iSNVs at random. In all, swine data subsets 1 and 2 each include 85 paired-time observations. Table S2 lists the identified iSNVs in the swine data, with further information on which paired-time observations were included in swine data subset 1 and which were included swine data subset 2. Figure 4A plots the frequency dynamics of the iSNVs in swine data subset 1. Figure 5B plots the frequency dynamics of the iSNVs in swine data subset 2.

**Figure 4.**
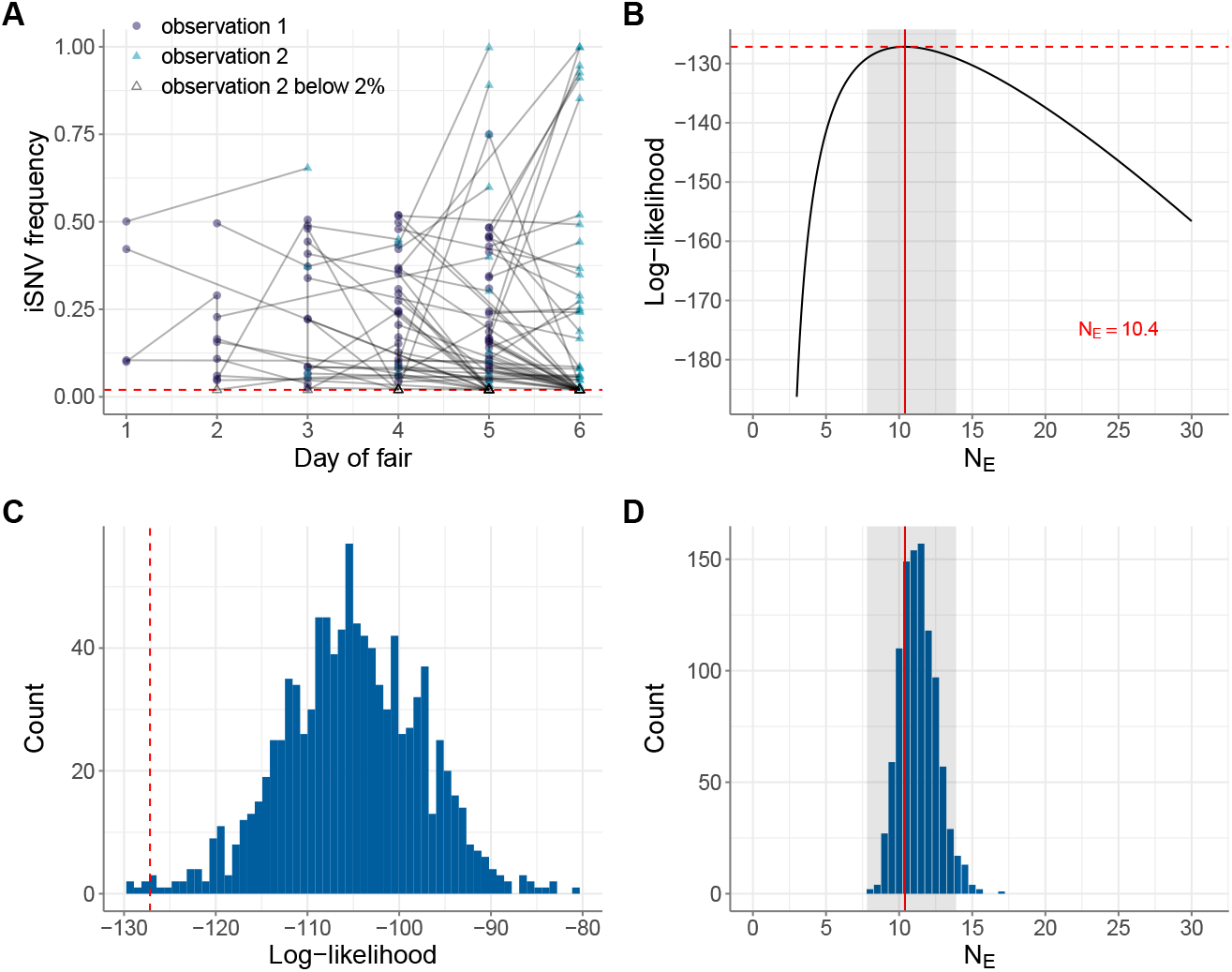
Estimates of within-host IAV effective population size from iSNVs present in swine data subset 1. (A) Allele frequency changes between the first observation time point and the second observation time point. Allele frequencies are plotted by day of county fair. The red dashed line shows the variant-calling threshold of 2%. Allele frequencies that fall under this threshold are shown at the threshold. (B) Calculated log-likelihood values across a range of effective population sizes. Solid red line shows the maximum likelihood estimate (MLE) of *N*_E_. Dashed red line shows the log-likelihood value for the MLE of *N*_E_. The shaded region shows the 95% confidence interval around the MLE of *N*_E_. (C) Calculated log-likelihood values for 1000 mock iSNV datasets that were generated by forward simulation of the Beta-with-Spikes model with *N*_E_ = 10.4 (blue histogram). The log-likelihood value at *N*_E_ = 10.4 calculated from swine data subset 1 is shown with a dashed red line. (D) Maximum likelihood estimates of *N*_E_ from 1000 simulated datasets, obtained using the Beta-with-Spikes model (blue histogram). The MLE of *N*_E_ = 10.4 from swine data subset 1 is shown with a solid red line. The shaded region shows the 95% confidence interval shown in panel B.

**Figure 5.**
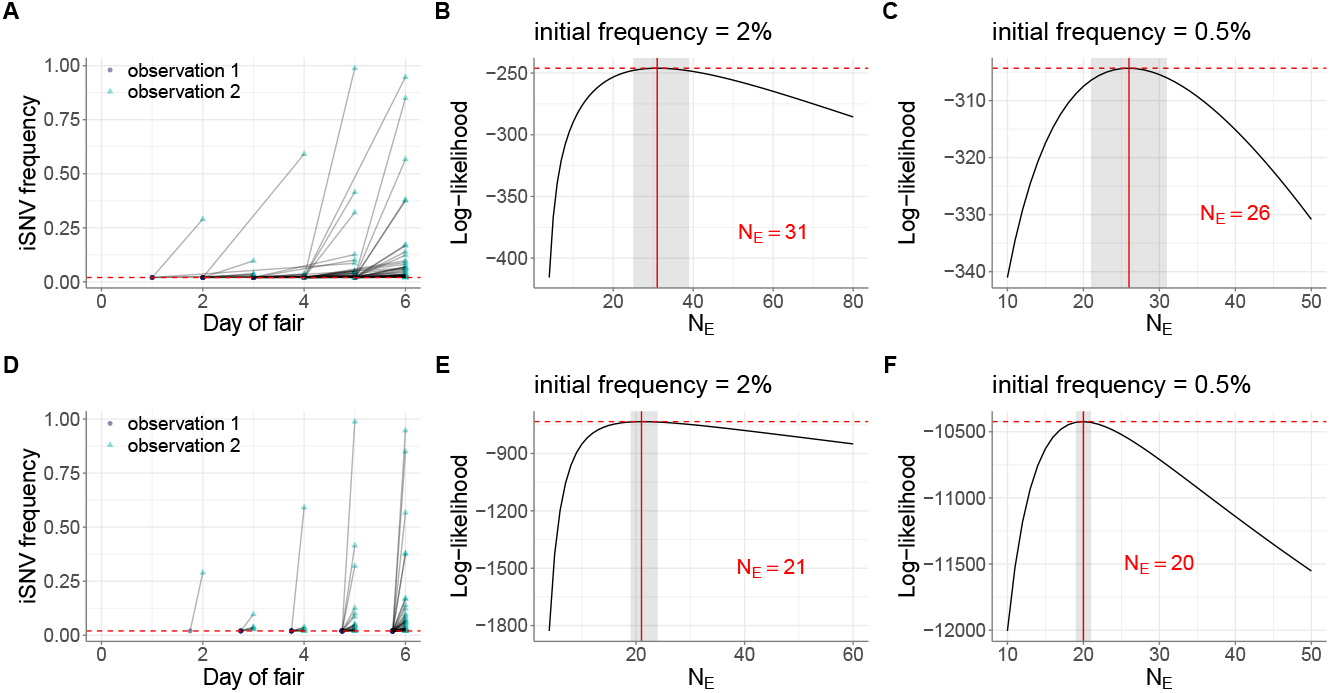
Estimates of within-host IAV effective population size from iSNVs present in swine data subset 2. (A) Allele frequencies at the first and second observation time points in swine data subset 2. As in Figure 4A, iSNV frequencies are plotted according to the day of the county fair and the dashed red line shows the variant-calling threshold of 2%. Below-the-variant-calling-threshold iSNV frequencies at the first time point are shown at 2%. (B) Calculated log-likelihood values across a range of effective population sizes when iSNVs at the first observation time point are assumed to be present at a frequency of 2%. (C) Calculated log-likelihood values across a range of effective population sizes when iSNVs at the first observation time point are assumed to be present at a frequency of 0.5%. (D) Allele frequencies from the paired-time observations in swine data subset 2, when the first time point is assumed to be a single viral generation prior to the second observation time point. As in Figure 5A, iSNV frequencies are plotted according to day of the county fair and the dashed red line shows the variant-calling threshold of 2%. Below-the-variant-calling-threshold iSNV frequencies at the first time point are shown at 2%. (E) Calculated log-likelihood values across a range of effective population sizes for the modified swine data subset 2 shown in panel (D). (F) As in panel (E), calculated log-likelihood values across a range of effective population sizes when iSNVs at the first time point are assumed to instead be present at a frequency of 0.5%. In panels (B), (C), (E), and (F), the solid red lines show the MLE of *N*_E_ and dashed red lines show the log-likelihood value for the MLE of *N*_E_. The shaded regions show the 95% confidence interval around the MLE of *N*_E_.

### Bioinformatic processing

We reanalyzed the sequence data that were made publicly available in McCrone et al. (2018) and VanInsberghe et al. (2024) in order to have a consistent analysis across human IAV and swine IAV datasets. Sequencing reads were first processed with fastp v.0.23.4 (Chen et al., 2018) to remove reads shorter than 60 nucleotides (nt) or with complexity less than 30%, trim adapters (with auto-detection) and 3’ poly X runs longer than 10 nt, and perform base correction in overlapped regions. Influenza genomes were assembled and sample-specific consensus sequences inferred using the Iterative Refinement Meta-Assembler (IRMA) FLU v.1.1.4 (Shepard et al., 2016). Assembled reads from each influenza segment were re-aligned to the sample-specific reference for that segment using Bowtie 2 v.2.5.4 (Langmead and Salzberg, 2012) with default settings (end-to-end mode). LoFreq (Wilm et al., 2012) was used to probabilistically realign reads and call variants relative to sample-specific consensus genomes. Snakemake v.7.32.4 (Mölder et al., 2021) was used for bioinformatic workflow management. Called variants were further filtered based on their estimated allele frequencies using Numpy v.2.0.1 (Harris et al., 2020) and Pandas v.2.2.2 (Pandas Development Team, 2020) in Python v.3.12.4 (Python Software Foundation, 2020). Figures S1 and S2 show the iSNV frequencies arrived at by our pipeline against those from the original analyses (McCrone et al., 2018; VanInsberghe et al., 2024).

### Estimation of effective population size *N*_E_

We estimate the effective population size of within-host human and swine IAV populations by separately interfacing the Beta-with-Spikes model, described above, with human data subsets 1 and 2 and swine data subsets 1 and 2. For both data subsets 1, we set the initial DAF at time *t*_0_ to have a mean *p*_0_ given by the observed initial iSNV frequency at the first time point of sampling and a variance given by *v*_0_ = *mp*_0_(1 − *p*_0_). We set the parameter *m* to 0.004 but our results were not sensitive to the exact value of *m* assumed. To quantify the probability of observing an allele at a given frequency at the second observation time point, we forward simulate equation (1) under a specified *N*_E_ from generation *t*_0_ to generation *t*_*k*_, where *k* denotes the number of viral generations between the first time point and the second time point in a set of paired-time iSNV observations. We assume that a viral generation is 6 hours long, based on experimental data (Dou et al., 2017; Einav et al., 2020) and parameter estimates from quantitative models fit to IAV kinetic data (Baccam et al., 2006). As such, the number of viral generations between a pair of iSNV observations is given by *k* = (24*/*6)*d* = 4*d*, where *d* is the number of days between the observed time points.

When the frequency of the focal allele at the second time point is above the variant-calling threshold, the probability of observing this data point is simply the probability given by the Beta-with-Spikes distribution evaluated at the observed frequency. When the frequency of the focal allele at the second time point falls below the variant-calling threshold, we calculate the probability of observing this data point by using the cumulative density function (cdf), evaluated at the variant-calling threshold. This cdf integrates the probability density function over the frequencies [0, *f*_*th*_), where *f*_*th*_ denotes the variant-calling threshold. The overall log-likelihood for a given *N*_E_ is given by the sum of the logs of the calculated probabilities across all paired-time observations in data subset 1. For data subsets 2, we set the initial DAF at time *t*_0_ to have a mean of *p*_0_ = *f*_*th*_ and again a variance of *v*_0_ = *mp*_0_(1 − *p*_0_), where we set parameter *m* to 0.004. The remaining analyses are analogous to those for data subsets 1.

### Assessing model misspecification

To determine whether observed changes in allele frequencies are consistent with a Wright-Fisher model, we generated mock datasets and applied the Beta-with-Spikes model to these mock data. The mock datasets were generated by setting initial iSNV frequencies at time *t*_0_ to those in data subset 1 and setting the number of viral generations for those iSNVs to match those in the empirical dataset. For each iSNV, we then forward-simulated the Beta-with-Spikes model to the second observation time point under the maximum likelihood estimate of *N*_E_ for data subset 1 and sampled an iSNV frequency from the simulated DAF. We generated 1000 of these mock datasets. For each of these mock datasets, we inferred *N*_E_ and kept track of the log-likelihood value that corresponds to the *N*_E_ maximum likelihood estimate. We then assessed whether the Beta-with-Spikes model could recover the *N*_E_ value that was used in the simulation of the mock data subsets 1, with log-likelihood values that were similar to those of the empirical data subset 1. Higher log-likelihood values for the mock datasets compared to the maximum log-likelihood value for the empirical data subset 1 would indicate that the assumed Wright-Fisher model may be misspecified and that an alternative evolutionary model may need to be considered to faithfully reproduce patterns of within-host IAV evolution.

## Results

### Within-host human IAV effective population sizes are small

Application of the Beta-with-Spikes model to human data subset 1 resulted in a maximum likelihood estimate of *N*_E_ = 41 viral particles (95% CI = [22,72]) (Figure 2B). This estimate of *N*_E_ is consistent with the *N*_E_ estimate of 32-72 viral particles from McCrone et al. (2020) that used a diffusion approximation. In both cases, the estimated *N*_E_ values are very small, underscoring the dominant role that genetic drift plays in the evolution of IAV populations within acutely-infected humans.

Human data subset 2, comprising pairs of sampling points where the iSNV is called only in the second time point, presents a greater challenge for inference. The issue is that, while the Beta-with-Spikes model easily handles iSNVs that start at an intermediate frequency and then are lost or fixed due to drift, for iSNVs that start rare or absent and then reach observable frequency, the inferred *N*_E_ is sensitive to the unobserved starting frequency of the iSNV. At one extreme, one can assume that the iSNV was just barely under the calling threshold frequency of 2% at the first sampled time point; such trajectories are shown in Figure 3A. This minimizes the inferred frequency change and maximizes the time over which that change occurred, and thus represents the minimum amount of drift (maximum *N*_E_) that is consistent with data. In human data subset 2, this assumption resulted in a maximum likelihood estimate of *N*_E_ = 335 viral particles (95% CI = [190, 610]) (Figure 3B). This estimate of *N*_E_ is considerably larger than the *N*_E_ ≈ 40 estimated from human data subset 1. Alternatively, the minimum amount of time over which the iSNV could have changed in frequency from below to above the limit of detection is a single viral generation. Figure 3D shows iSNV trajectories assuming that they start from just below 2% frequency in the generation before they are observed. Note that while the magnitude of frequency change is the same as in Figure 3A, it occurs in much less time, implying a smaller *N*_E_. Indeed, this assumption yielded a MLE of *N*_E_ = 52 (95% CI = [32-75]) (Figure 3E), consistent with the estimate from human data subset 1. We also tested the effect of reducing the assumed iSNV starting frequency from 2% to 0.5%, increasing the magnitude of inferred frequency change. As expected, this reduced the inferred *N*_E_ values relative to those obtained with a 2% starting frequency. Assuming that the iSNVs were at frequency 0.5% in the first sampling point yielded a MLE of *N*_E_ = 176 (95% CI = [126, 246]) (Figure 3C). Assuming that they were present at a frequency of 0.5% in the viral generation before they were observed yielded a MLE of *N*_E_ = 35 (95% CI = [32, 37]) (Figure 3F). Human data subset 2 is thus consistent with a wide range of possible values for *N*_E_, including the *N*_E_ ≈ 40 estimated from human data subset 1. Together, the two data subsets point to effective population sizes of under 100 for seasonal IAV populations within acutely infected humans.

### iSNV frequency changes in human infections are consistent with a Wright-Fisher model of evolution

To determine whether the observed iSNV frequency changes in the human IAV infections were consistent with a basic Wright-Fisher model with a small effective population size, we quantitatively analyzed our mock human data subsets. As expected, application of the Beta-with-Spikes model to these simulated datasets resulted in recovery of *N*_E_ estimates close to the *N*_E_ = 41 value that was used during their generation (Figure 2D). Moreover, Figure 2C shows the log-likelihood values of these simulated datasets, evaluated at the MLE of each dataset’s *N*_E_, along with that of the empirical human data subset 1. The log-likelihood value of −30.7 for human data subset 1 falls squarely within the distribution of log-likelihoods of the mock datasets. This indicates that human data subset 1 is consistent with a Wright-Fisher model of evolution. If it were inconsistent with this model, we would expect that the log-likelihood values from the simulated datasets to be exceed those from the empirical human data subset 1.

### iSNV frequency dynamics in within-host swine IAV populations also point to the importance of genetic drift, but suggest discrepancy from a Wright-Fisher model of evolution

Figure 4 shows estimates of *N*_E_ based on the application of the Beta-with-Spikes model on swine data subset 1. The MLE for this dataset is *N*_E_ = 10 (95% CI = [8, 14]) (Figure 4B), pointing towards genetic drift being a dominant driver of within-host IAV evolution in pigs. Application of this approach to the mock swine datasets resulted in successful recovery of this small *N*_E_ (Figure 4D). Interestingly, however, the log-likelihood at the MLE of *N*_E_ = 10 for empirical swine data subset 1 was lower than all but 6 mock datasets (0.6%) that were simulated under the Beta-with-Spikes Wright-Fisher model. This indicates that the distribution of iSNV frequency changes observed in swine data subset 1 is not consistent with a classic Wright-Fisher model. That is, these results allow us to reject the hypothesis that iSNV frequency changes in within-host swine IAV populations are solely governed by a simple Wright-Fisher process of genetic drift.

Finally, we applied the Beta-with-Spikes model to swine data subset 2. Figure 5 shows these results, with figure panels analogous to those shown for Figure 3 for human data subset 2. In the case of swine data subset 2, we estimated *N*_E_ to be 31 viral particles under the assumption that the iSNVs in this dataset were present at 2% at the first observation time point (Figure 5B). This estimate exceeded that for swine data subset 1, similarly to how the analogous estimate on human data subset 2 exceeded that for human data subset 1. As expected, resetting the iSNV frequencies at the first observation time points to a lower threshold of 0.5% lowered the *N*_E_ estimate (Figure 5C), but the estimate still exceeded that from swine data subset 1. Finally, assuming that the first observation time points instead fell a single viral generation before the second observation time points yielded *N*_E_ estimates of 21 and 20 for iSNV frequencies set to 2% and 0.5% at these first time points, respectively (Figures 5E,F).

## Discussion

Several previous studies have underscored the prominent role that genetic drift plays in shaping the evolutionary dynamics of influenza A virus populations in acutely infected individuals (Dinis et al., 2016; Debbink et al., 2017; McCrone et al., 2018, 2020; VanInsberghe et al., 2024; Bendall et al., 2024). Here, we reanalyzed two previously published datasets (one human IAV dataset and one swine IAV dataset) to quantify the extent of genetic drift acting on within-host IAV populations. Consistent with a previous study that estimated the effective viral population size from the same set of acutely infected humans to be *N*_E_ = 32 *−* 72 (McCrone et al., 2020), we here similarly found evidence of a very small *N*_E_ of around 40 viral particles. The consistency of these findings is reassuring, given that the previous study used a diffusion approximation that can break down at small *N*_E_ values. Our estimate was instead derived from a recently developed Beta-with-Spikes model that has been shown to provide accurate estimates for both small and large *N*_E_ values (Tataru et al., 2015). There are several other differences between the approach taken by McCrone et al. (2020) and the one we used here. McCrone et al. (2020) used all of the iSNV frequency data at once and jointly estimated *N*_E_ and the mutation rate *µ*. We instead partitioned the data into two subsets. The first subset included the iSNVs that were detected above the variant-calling threshold at the first observation time point and the second included the iSNVs where iSNVs were identified above the variant-calling threshold at the second observation time point but were not called at the first observation time point. This partitioning allows us to avoid estimation of the mutation rate, which should impact not only sites with observed iSNVs but also those that do not show evidence of polymorphism. Our approach and the approach taken by McCrone et al. (2020) are complementary, and the consistency of our *N*_E_ estimates underscores the robustness of the results. Our estimates of *N*_E_ in the swine IAV dataset are to our knowledge the first such estimates for within-host IAV infections in acutely infected pigs.

Recent work by Bendall et al. (2024) that analyzed the evolutionary dynamics of seasonal IAV populations within acute human infections also estimated *N*_E_. The estimates of *N*_E_ = 176 − 284 arrived at in this study are considerably higher than those estimated with the previous human IAV dataset from the same lab (Lauring) (McCrone et al., 2018, 2020) that we also examined here. We do not know the reason for the difference in *N*_E_ estimates between these two human IAV datasets. One possibility, and our favored hypothesis, is that this difference has to do with the difference in variant-calling threshold applied to the datasets. In McCrone et al. (2018), a 2% threshold was applied, whereas in Bendall et al. (2024), a 0.5% threshold was applied. While the choice of threshold should not impact point estimates of *N*_E_ (just the breadth of the confidence intervals), spurious variants would tend to bias *N*_E_ upwards. While spurious variants occur more commonly at lower variant-calling thresholds, Bendall et al. (2024) did perform a benchmarking analysis to determine how low they could go with their variant-calling threshold. If their *N*_E_ estimate was inflated due to the presence of spurious variants at this low 0.5% variant-calling threshold, they may also be over-detecting positive selection. This is because the approach used by Bendall et al. (2024) sequentially estimates *N*_E_ and then selection coefficients. At large values of *N*_E_, dramatic changes in iSNV frequencies have to invoke selection, as genetic drift cannot explain these changes. If *N*_E_ is actually smaller than estimated, then a subset of the dramatic changes in iSNV frequencies will therefore no longer need to invoke selection, as genetic drift could explain the changes. Overestimation of *N*_E_ due to a too-low variant-calling threshold might therefore explain why this previous work identified a number of synonymous iSNVs to be under positive selection. In any case, it would be interesting to gauge the robustness of the *N*_E_ estimate from the Bendall et al. (2024) dataset under a higher variant-calling threshold of 2%.

Beyond estimation of within-host effective viral population sizes, we here simulated mock datasets and used these datasets to help us address the question of whether the empirical datasets are consistent with the classic Wright-Fisher model of evolution. Through this analysis, we found that the iSNV frequency changes observed in the human IAV dataset were consistent with the Wright-Fisher model. In contrast, we found that the evolutionary dynamics observed in the swine IAV dataset could not be faithfully reproduced by the classic Wright-Fisher model. This result is critical and indicates that future studies need to focus on determining what other processes might be contributing to within-host IAV evolution. Other processes may include highly skewed offspring distributions as well as spatial compartmentalization. Identification of these additional processes will allow us to not only better dissect the roles of different evolutionary processes that occur within hosts but also to better understand the processes that guide the path from mutation generation to viral spread at the population level.

## Data Availability

All inference code will be made available on GitHub at: https://github.com/koellelab/IAV_beta_with_spikes_Ne

