## Supplemental Material for "Genetic drift acts strongly on within-host influenza virus populations during acute infection but does not act alone"

**Yike Teresa Shi[1], Michael A. Martin[2], Daniel Weissman[3], Katia Koelle[1,4,*]**

**1 Department of Biology, Emory University, Atlanta, GA, USA**
**2 Department of Pathology, Johns Hopkins School of Medicine, Baltimore, MD, USA**
**3 Department of Physics, Emory University, Atlanta, GA, USA**
**4 Emory Center of Excellence for Influenza Research and Response (CEIRR), Atlanta GA, USA**

**[*]**

### Supplemental Figures

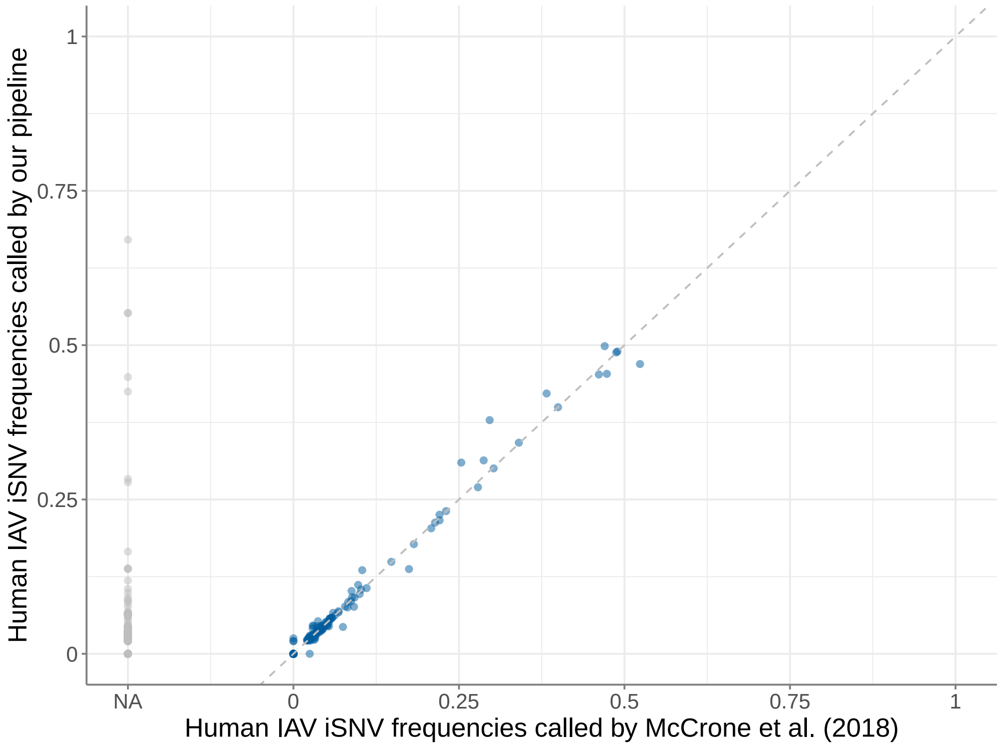


**Figure S1.** Comparison between iSNV frequencies called in the original analysis of McCrone et al. (2018) and those called by our pipeline (detailed in the Methods section).

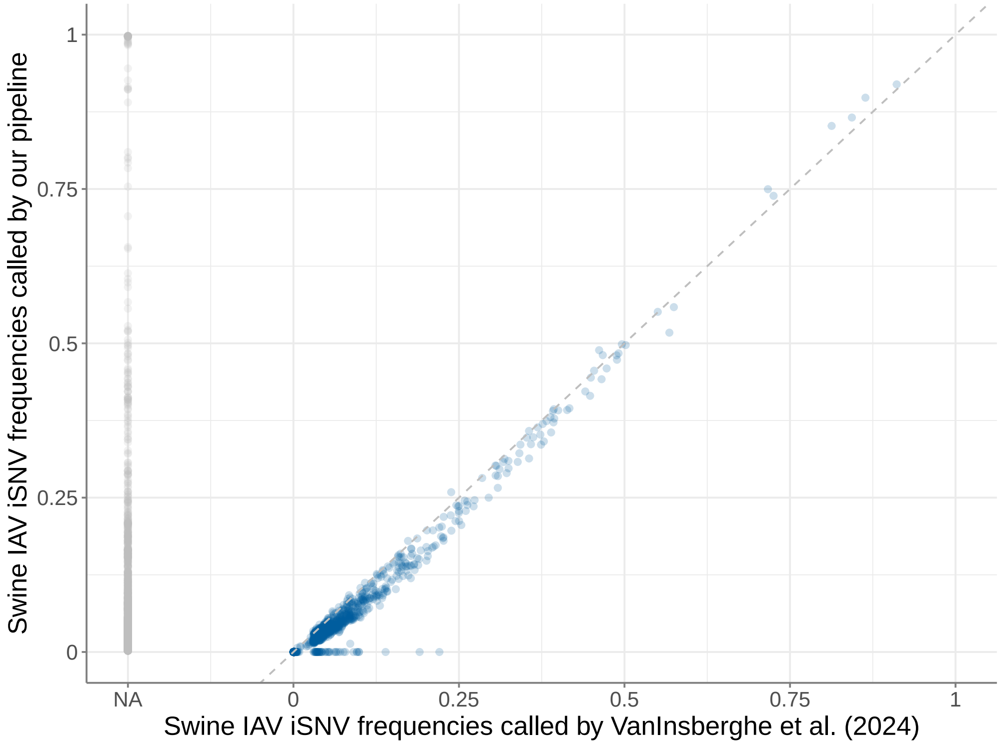


**Figure S2.** Comparison between iSNV frequencies called in the original analysis of VanInsberghe et al. (2024) and those called by our pipeline (detailed in the Methods section).
